# Hippocampal-Prefrontal cortex network dynamics predict performance during retrieval in a context-guided object memory task

**DOI:** 10.1101/2021.11.28.470274

**Authors:** JF Morici, NV Weisstaub, CL Zold

## Abstract

Remembering life episodes is a complex process that requires the interaction between multiple brain areas. It is thought that contextual information provided by the hippocampus (HPC) can trigger the recall of a past event through the activation of medial prefrontal cortex (mPFC) neuronal ensembles, but the underlying mechanisms remain poorly understood. Indeed, little is known about how the vHPC and mPFC are coordinated during a contextual-guided recall of an object recognition memory. To address this, we performed electrophysiological recordings in behaving rats during the retrieval phase of the object-in-context memory task (OIC). Coherence, phase locking and theta amplitude correlation analysis showed an increase in vHPC-mPFC LFP synchronization in the theta range when animals explore contextually mismatched objects. Moreover, we identified ensembles of putative pyramidal cells in the mPFC that encode specific object-context associations. Interestingly, the increase of vHPC-mPFC synchronization during exploration of the contextually mismatched object and the preference of mPFC incongruent object neurons predicts the animals’ performance during the resolution of the OIC task. Altogether, these results identify changes in vHPC-mPFC synchronization and mPFC ensembles encoding specific object-context associations likely involved in the recall of past events.

## Introduction

Revisiting in our minds events that have occurred in the past is critical for survival. This cognitive function is called episodic memory and integrates information of past events regarding “what”, “when” and “where” it happened (Templer and Hampton, 2013). The hippocampus has been postulated as a key structure in encoding and retrieval of episodic memories due to its ability to codify contextual information (Zemla and Basu, 2017) as well as spatial navigation (Eichenbaum, 2017; Moser et al., 2017). Particularly, during recall, contextual information should impact on the activity of neocortical areas, also identified as nodes involved in episodic memory retrieval (Eichenbaum, 2017b; Wirt & Hyman, 2019). Especially, the Prefrontal Cortex is postulated to exert a controlling role over hippocampus dependent memories by establishing relationships among memories and monitoring retrieval (Eichenbaum, 2017b; Zhang et al., 2018). Imaging studies as well as mPFC lesions in humans strongly suggest that this structure is recruited specifically under conditions of memory interference or distractions (Tomita et al., 1999; Eichenbaum, 2017c). Evenmore, tracing studies have shown a robust direct, monosynaptic, ipsilateral projection from the vHPC to the mPFC (Hoover and Vertes, 2007) as well as indirect connections through the Reuniens nucleus of the thalamus (Varela et al., 2013; Hoover and Vertes, 2011). Then it is hypothesized that, during retrieval, the vHPC sends contextual information that engages previously formed contextual rules in the mPFC to support memory recall and to inhibit interference from less relevant memory traces (Eichenbaum 2016, Eichenbaum, 2017c). However,the mechanisms underlying this vHPC-mPFC interaction remain unknown.

Cognition is associated with an oscillatory pattern of neural activity in the brain that involves several frequency bands simultaneously. Slow rhythms allow information sharing by synchronizing distant brain areas. Particularly, theta rhythms in the HPC have been linked with exploratory behavior (Buzsáki and Moser, 2013; Buzsáki and Tingley, 2018), memory formation, and recall (Hasselmo and Howard Eichenbaum, 2005; Boyce et al., 2016; Lopes-dos-Santos et al., 2018). In the case of the mPFC, theta rhythms had been associated with attention processes (O’Neill et al., 2013), and memory recall (Marko et al., 2019; Riddle et al., 2020). Furthermore, it has been shown that coordination of theta oscillations between HPC and the mPFC increases during free exploration (Siapas et al., 2005), anxiety-related behaviors (Adhikari et al., 2010; Adhikari et al., 2011), decision making (Tang et al., 2021) as well as mnemonic processes that are spatial/contextually guided and rewarded (Benchenane et al., 2010; Place et al., 2016. Also, single-cell activity in the mPFC is also modulated by theta synchronization in a task-specific manner (Adhikari et al., 2010). However, little is known about the interplay between the vHPC and the mPFC during retrieval of episodic-like memories, and how the transference of information between these structures depends on contextual information.

Novel object recognition tasks are non-rewarded paradigms based on the natural tendency of rodents to explore novel objects. These tasks are used to investigate episodic-like memory in rodents (Morici et al 2015; Barker and Warburton, 2020b). Some versions require contextually-guided recall to recognize object-context matches. This is the case of the Object-in-context (OIC) task, where animals need to recognize which of two familiar objects is contextually mismatched (Barker and Warburton, 2020). Both the mPFC and the vHPC, as well as the interaction between them, are required for the resolution of this task. In this work, we analyzed the functional connectivity between the mPFC and the vHPC in rats by simultaneously recording neural activity from the vHPC and mPFC in freely moving rats during the resolution of the OIC task. We found an increase in theta oscillation power in the mPFC and vHPC as well as an increased synchronization between both structures during the retrieval phase, as assessed by coherence analysis and correlation analysis. The increase in coherence is maximum when the animals are exploring the contextually incongruent object and it correlates with recall performance during the task. We also identified cells in the mPFC that respond differently to the object in context combinations and whose preference correlates with the animals’ recall performance.

## Materials and Methods

*Ethical Statement:* All experimental procedures were in accordance with the institutional Animal Care and Use Committee (School of Medicine, University of Buenos Aires, ASP # 49527/15, and University of Favaloro #DCT0205-18) and government regulations (SENASAARS617.2002). All efforts were made to minimize the number of animals used and their suffering.

*Subjects:* A total of 22 Wistar male rats were used: 9 for electrophysiological recordings and 13 for pharmacological experiments. Two animals were excluded due to poor electrophysiological signal-to-noise ratio. One more animal was excluded due to electrode misplacement leaving a total of 6 animals for electrophysiological recordings. No animals were excluded from the pharmacological experiments.

At the time of the surgery, animals were >P60, weighted 250-300 gr, and were housed in groups. Animals used for chronic electrophysiological recordings were single housed after surgery. Rats were kept with water and food ad libitum under a 12 hr light/dark cycle (lights on at 7:00 A.M.) at a constant temperature of 21-23°C. Experiments took place during the light phase of the cycle (between 10:00 A.M. and 5:00 P.M.) in quiet rooms with dim light.

### Surgical procedures

#### For electrophysiological recordings

Animals were anesthetized with isoflurane, treated with local anesthetic (bupivacaine hydrochlorate solution, 5% wv/v, Durocaine, AstraZeneca S.A., Argentina, 0.2-0.3ml s.c.) in the scalp and pressure points, and secured within a stereotaxic frame. The temperature was maintained using a heating pad. The skull was exposed and adjusted to place bregma and lambda on the same horizontal plane. Then, small metal screws were placed as anchors for the implant. Craniotomies were drilled to place the electrodes within the prefrontal cortex (mPFC, +3.2 mm anterior, +/- 1.2 mm lateral from bregma, and -2 mm from brain surface) and the ventral hippocampus (vHPC, -5.75 mm anterior, +/- 4.6 mm lateral from bregma, and -6.3 mm from brain surface), with an angle of 10° and 350° respectively (**Figure 2A**). The array designed to record within the mPFC consisted of four tetrodes (made of 20-μm HML-insulated tungsten wire, final impedance adjusted by gold plating to 150-250 kOhms) separated by 250 μm in a square shape (**Figure 2A**). For the vHPC, two bipolar electrodes separated by 250 μm were used (50-μm Formvar-insulated Nickel Chrome wire) (**Figure 2A**). A screw located on the occipital bone was used as the ground for the electrophysiological recordings. Sterile petroleum ointment was applied to the craniotomy, and dental cement was used to anchor the electrode assemblies to the skull. Animals were then treated with antibiotics (gentamicin, 1 mg/ml), a non-steroidal anti-inflammatory agent (flunixin, 4 mg/ml), and kept under careful observation until they were awake. Animals were allowed to recover in their home cage for at least 1 week and received dietary supplements until their weight was stabilized before conducting the behavioral experiments.

**Figure 1:**
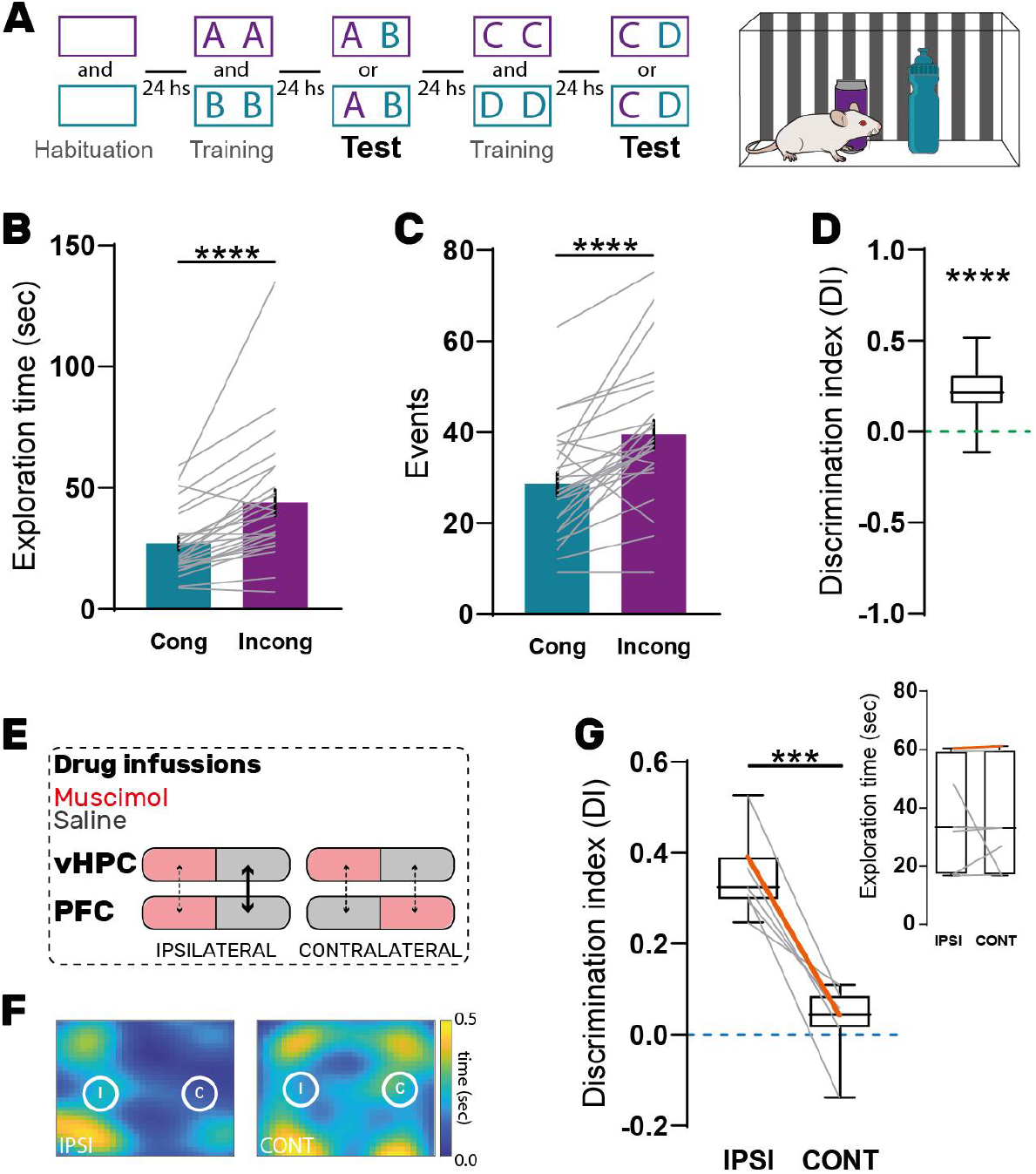
Ventral hippocampus and Prefrontal cortex are functionally connected during the resolution of the Object-in-Context task. (A) Each Object-in-Context task (OIC) block is a 5 day procedure. Each column represents a day. Animals were first habituated to two different contexts and then trained to two object context associations. 24hs after the training session animals are reexposed to one of the contexts and copies of both previously presented objects. (B) Mean exploration time and (C) number of exploration events for the Congruent (blue bar) and Incongruent object (purple bar) obtained during test sessions from 5 rats. Paired t test, N=21, **** p=0.002, t_B_=4.58, t_C_=4.52. Graphs represent mean ± SEM (D) Discrimination Index (DI) calculated for all sessions. One-sample t test against zero, ****p<0.0001, t=8.48. The graph represents mean ± max and min. (E) Infusions procedure for pharmacological disconnection experiment. Muscimol (Red) or Saline (Grey) were infused ipsilaterally (IPSI) or contralaterally (CONT) in the same animal 15 minutes before each test session. The same animal received 4 injections in a pseudorandomly manner for each test session. (F) Heat-map representing the centroid of an example animal in IPSI and CONT conditions. Note that animals explore more the incongruent object (left) in the IPSI condition, while they equally explore both objects if muscimol is injected contralaterally. (G) Discrimination Index calculated in IPSI and CONT conditions. Paired t test, N=7 per group, ***p=0.003, t=7.62. Inset: Total object exploration time. Paired t test, p=0.59, t=0.56. Red line indicates the example shown in F. Graphs represent median, 25-75 quartiles and range.

**Figure 2:**
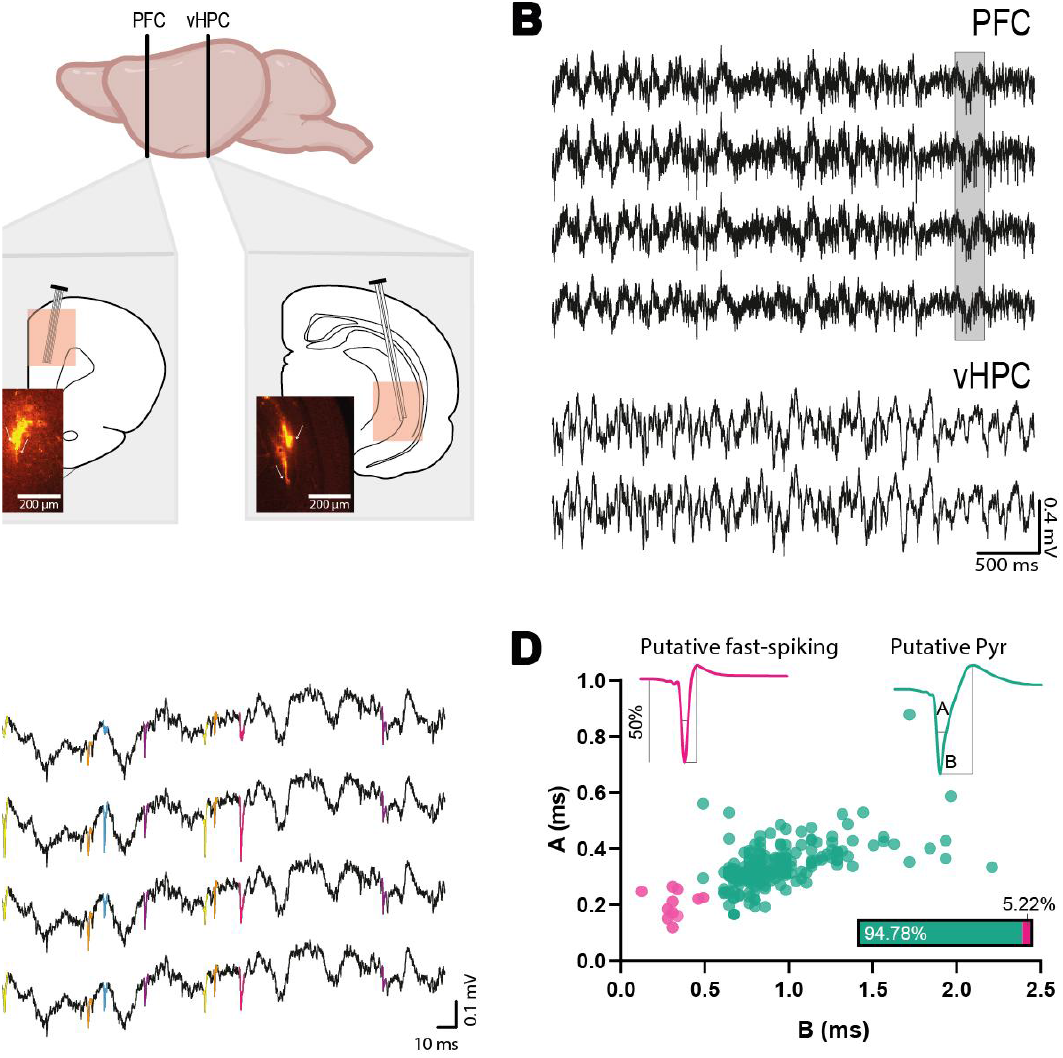
Electrophysiological recordings obtained during the resolution of the OIC task. (A) Electrode location. Dye-staining of the trace of mPFC tetrodes and vHPC stereodes. (B) Raw LFP signals from one mPFC tetrode and a vHPC stereode simultaneously recorded during the test phase of the task. (C) Example of mPFC raw signal showing spikes colored according to their clustering into separate single-units. (D) mPFC pyramidal cell and interneuron classification based on their electrophysiological properties. Examples of isolated unit waveforms classified as interneurons (pink) and pyramidal (green) cells. The parameters used for the classification were: A, half-amplitude duration; and B, trough to peak time. The parameters A and B were plotted for each cell, and two clear clusters were formed and used to sort neurons into putative pyramidal and interneurons.

#### Pharmacological disconnection experiments

Animals were deeply anesthetized with ketamine (Inducmina, 80 mg/kg i.p.) and xylazine (Richmond, 8 mg/kg i.p) and placed in a stereotaxic frame. The skull was exposed and adjusted to place bregma and lambda on the same horizontal plane. After small burr holes were drilled, two pairs of guide cannulae (22 G) were implanted bilaterally into the mPFC and vHPC (same coordinates mentioned above). At the end of the surgery and until the infusions were made, dummy injectors (30 G) were inserted into the guide cannula to preclude their obstruction. To confirm correct cannulae placement, 24 hrs after the end of the behavioral experiments, animals were infused with 1 ml of methylene blue through the dummy cannulae, deeply anesthetized, and transcardially perfused. Histological localization of the infusion sites were established using a stereo microscope (see Supp **Figure 2B** and **3C**).

**Figure 3:**
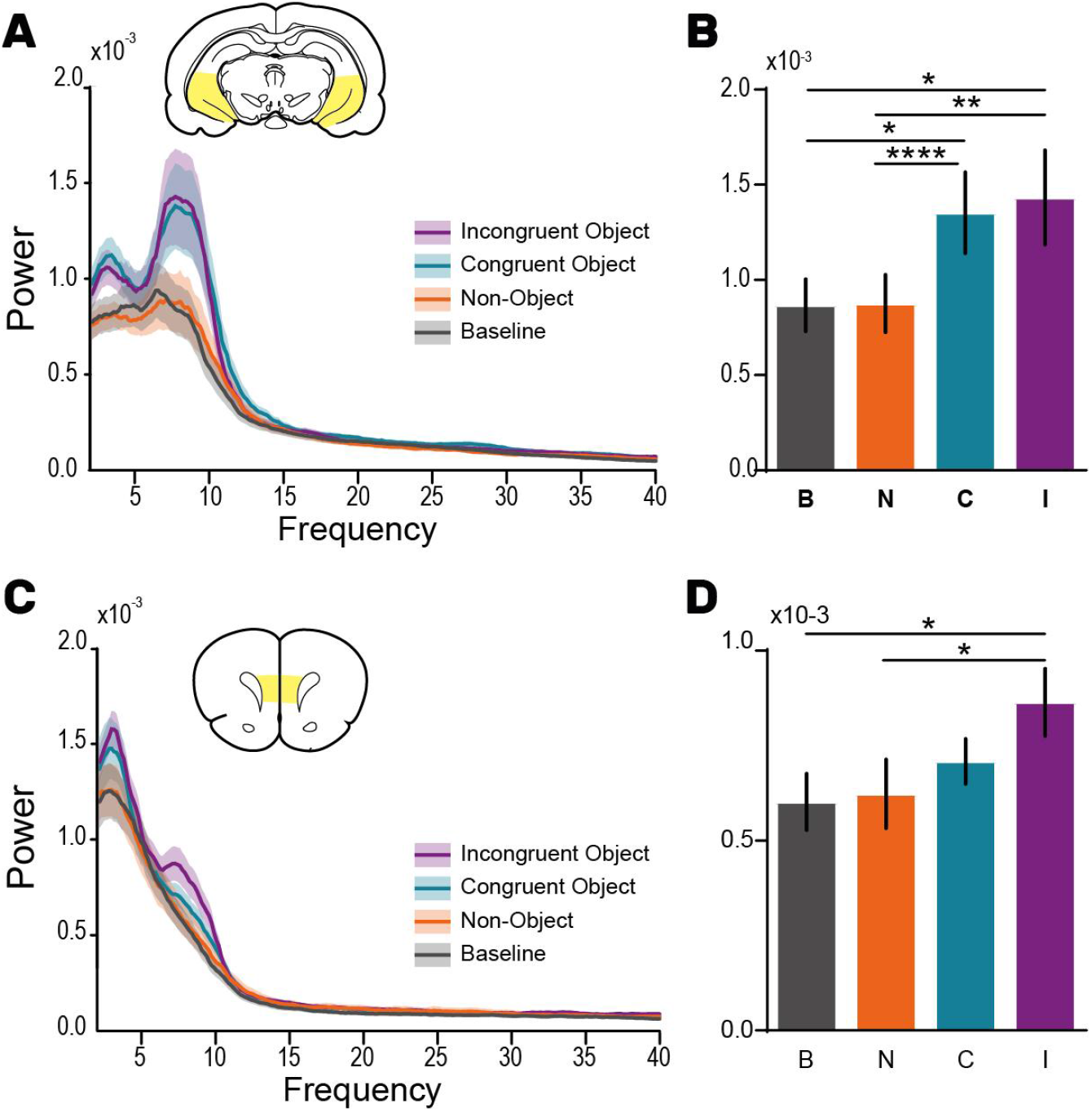
Theta oscillations are increased in the vHP and mPFC during object exploration. Power auto-spectrum of ventral hippocampal (A) and prefrontal cortex (C) LFP signals. Power spectral density analysis was performed using LFP segments recorded during baseline condition (Grey) and during the test session separating Non-Object (Orange), Congruent Object (blue) and Incongruent Object (purple) events. (B) Quantification of theta power in the vHPC for all conditions. Repeated measures One-way ANOVA, N=21, **p=0.002, F=7.44. Bonferroni’s post hoc, *p_C-B_=0.042, *p_I-B_=0.049, **p_I-N_=0.002, ****p_C-N_<0.0001. (D) Quantification of theta power in the mPFC. Repeated measures One-way ANOVA, N=21, p=0.01, F=4.02. Bonferroni’s post-hoc test, *p_I-B_=0.01, *p_I-N_=0.03. Bars represent mean ±SEM.

### Pharmacological disconnection intervention

In the test session, the dummy injector was removed and a 30 G injection cannula protruding1 mm below the guide cannula was inserted. The injection cannula was connected to a 10 ml Hamilton syringe. All animals were injected in four sites (mPFC and vHPC on both hemispheres) 15min before the test session. Animals received contralateral or ipsilateral infusions of Muscimol (MUS, 0.1 mg/ml, 1 µl per injection site, Alamone Lab, Cat #: M-240) and Vehicle (VEH, Saline 1µl) in a random manner. For instance, animals in the CONT group received MUS in the right mPFC and in the left vHPC (see Fig 1E).

### Behavioral paradigm

#### Apparatus

We used two differently shaped acrylic mazes (50 cm wide X 50 cm length X 39 cm height and 60 cm wide X 40 cm length X 50 cm height arenas). The floor and walls were white with different visual clues. In order to change the context each week, we designed removable walls with visual cues of different colors and shapes. Duplicate copies of objects made from plastic, glass, and aluminum were used. Objects were thoroughly cleaned between phases and randomly assigned to the different phases of the experiments. The heights of the objects ranged from 10 to 20 cm and they varied with respect to their visual and tactile qualities. All objects were affixed to the floor of the apparatus with an odorless reusable adhesive to prevent them from being displaced during each session. The objects were always located along the central line of the maze, away from the walls and equidistant from each other. As far as we could determine, the objects had no natural relevance for the animals and they were never associated with any reinforcement. The objects, floor, and walls were cleaned with ethanol 50% between sessions. Exploration of an object was defined as directing the nose to the object at a distance of <2 cm and/or touching it with the nose. Turning around or sitting on the object was not considered exploratory behavior.

#### Object-in-Context task (OIC)

This task is a three-trial procedure that allows evaluation of the congruency between the context and the object (Eacott, 2004. Bekinschtein et al., 2013). During the training phase, animals were sequentially exposed for 30 minutes to two different contexts each containing a different pair of identical objects. These presentations were separated by 1 hr. During the test phase, carried out 24 hrs after the last presentation, a new copy of each object used before is presented in one of the contexts (pseudorandomly assigned). Thus, one of the objects is presented in an ‘incongruent’ context (incongruent object), while the other is presented in a ‘congruent’ one (congruent object). In this task, novelty comes from a novel combination of an object and a context, and exploration is driven by retrieval of a particular ‘what’ and ‘which’ context conjunctive representation (Eacott, 2004). This session was 10 minutes long. To increase the number of recorded test sessions, we modified the task in order to increase the number of exposures in the following way: We divided the task into blocks. Each block involved a pair of distinct and different contexts, four pairs of objects and lasted 5 days. The objects were used in two pair sets (e.g: A & B) and were randomly chosen and assigned to each context in a balanced way across subjects. The selection of the context used in the test sessions was also random and balanced across the experiment. The basic procedure was: Day 1 - Habituation to Context 1 and 2, Day 2 - Training phase 1: novel pairs of objects A or B were presented in Context 1 or 2, Day 3 - Test session 1 (a copy of objects A and B were presented in either Context 1 or 2), Day 4 - Training phase 2 novel objects C or D were presented in Context 1 or 2, Day 5 - Test session 2 (copies of objects C and D objects were presented in either Context 1 or 2. At the end of Test session 2, animals were allowed to rest for 2 days before a different block was started (**Figure 1A**). Animals were then habituated to a novel set of contexts and presented with novel pairs of objects as previously described. All behavioral sessions were video recorded using a digital camera (Genius, FaceCam 1000x). Also, object exploratory events were online recorded and synchronized with the electrophysiological recordings by sending TTL signals to the amplifier through an Arduino. Based on the exploratory time, a discrimination Index (DI) was calculated as t_Incongruent_– t_congruent_/total exploration. We referred to the objects in the text as congruent and incongruent for descriptive purposes

#### Novel Object Recognition task (NOR)

We adapted the spontaneous novel object recognition task (Barker et al., 2007) used to evaluate recognition memories, as we did with the OIC task. Briefly, animals were exposed to 1 block of 2 consecutive trials. The block started with a single habituation session of 10 minutes to the context. On day 2 rats were exposed to a pair of novel identical objects for 30 minutes. On day 3 (test session 1) animals were re-introduced to the context used in the training phase and were allowed to explore the familiar object together with a novel one never presented before. On day 4, animals were reintroduced to the familiar context with a novel pair of objects and twenty-four hours later the second test session was conducted. All behavioral sessions were video recorded using a digital camera (Genius, QCam 6000). Exploration time was analyzed offline and based in the exploratory performance of the animals a Discrimination Index (DI) was calculated as t_novel_– t_familiar_/total exploratory time.

### Electrophysiology

Neural activity was recorded using a 32-channel Cheetah setup (Neuralynx). Neural signals were digitized, bandpass filtered (0.1 - 6000 Hz), and continuously sampled at 32 kHz. mPFC tetrodes were attached to a micromanipulator that allowed us to move them daily. Events in the behavioral task were simultaneously recorded by the acquisition system.

#### LFP Spectral Analysis

Since all wires within each brain structure showed very similar LFP signals, only one wire from the mPFC and vHPC was used hereon for the spectral analysis. LFP segments recorded during events of either congruent or incongruent object exploration were detrended, digitally-filtered (2-625Hz, plus notch filter) and concatenated using a Hamming window. Only exploration events longer than the median exploration time was considered for the analysis. The rate and duration of object exploration were higher for the incongruent object than for the congruent one. Thus, we randomly selected incongruent object exploration events to match the length of the congruent exploration time. For the non-object exploration events, we randomly selected periods of time when animals were in the task but not exploring the objects until reaching the duration of the less explored object time. The same procedure was applied for the baseline condition using signals acquired while animals were in their homecage before the test session. The power spectral (PSD), cross power spectral (cross-PSD), and coherence analyses were performed using the *coherencyc* function in the Chronux toolbox (http://chronux.org/). For PSD, cross-PSD, and coherence comparisons between conditions, we extracted each value at 7.5 Hz. For the coherence perievent spectrograms we used the *wcoherence* function from the Matlab signal processing toolbox. The coherence between the mPFC and the vHPC as calculated using signal segments of 4 seconds centered around the beginning and the end of the objects explorations events. This signal was then analyzed using 500 ms windows with 70% overlap and a mean matrix (Frequency x Time) was calculated for each session. The final arrays, used to construct the figures presented here, were the mean of all the arrays resulting from each session in both conditions (Congruent and Incongruent). The coherence max values from these spectrograms were extracted at the same frequency studied in the previous analysis (7-8 Hz) for the first and last 500ms of the object exploration events (ON and OFF perievent coherence spectrograms respectively). The same procedure was used to construct the coherence perievent spectrograms across test sessions, but only the coherence at the start of the object exploration events were analyzed. For each session, the total number of exploration events for each object were equally divided into three segments and the mean time x frequency matrices were constructed. For this analysis, only sessions with at least 5 exploration events were used.

#### LFP Amplitude Cross-correlation Analysis

We performed a vHPC-mPFC amplitude correlation as it was described elsewhere (Adhikari et al., 2010). First, we extracted mPFC and vHPC LFP signals during Congruent and Incongruent objects exploration events as described above. LFP signals were digitally filtered between 6 to 10Hz and a Hilbert transform was applied. The absolute value of the instantaneous Hilbert amplitude was used to compute a cross-correlation between the vHPC and mPFC theta oscillations amplitude using a lag of +/-200 ms for each object condition (**Figure 5H**). We also compared the area under the curve of the theta oscillation envelope of the object exploration events to the average z-score throughout the session (**Figure S4A**).

**Figure 4:**
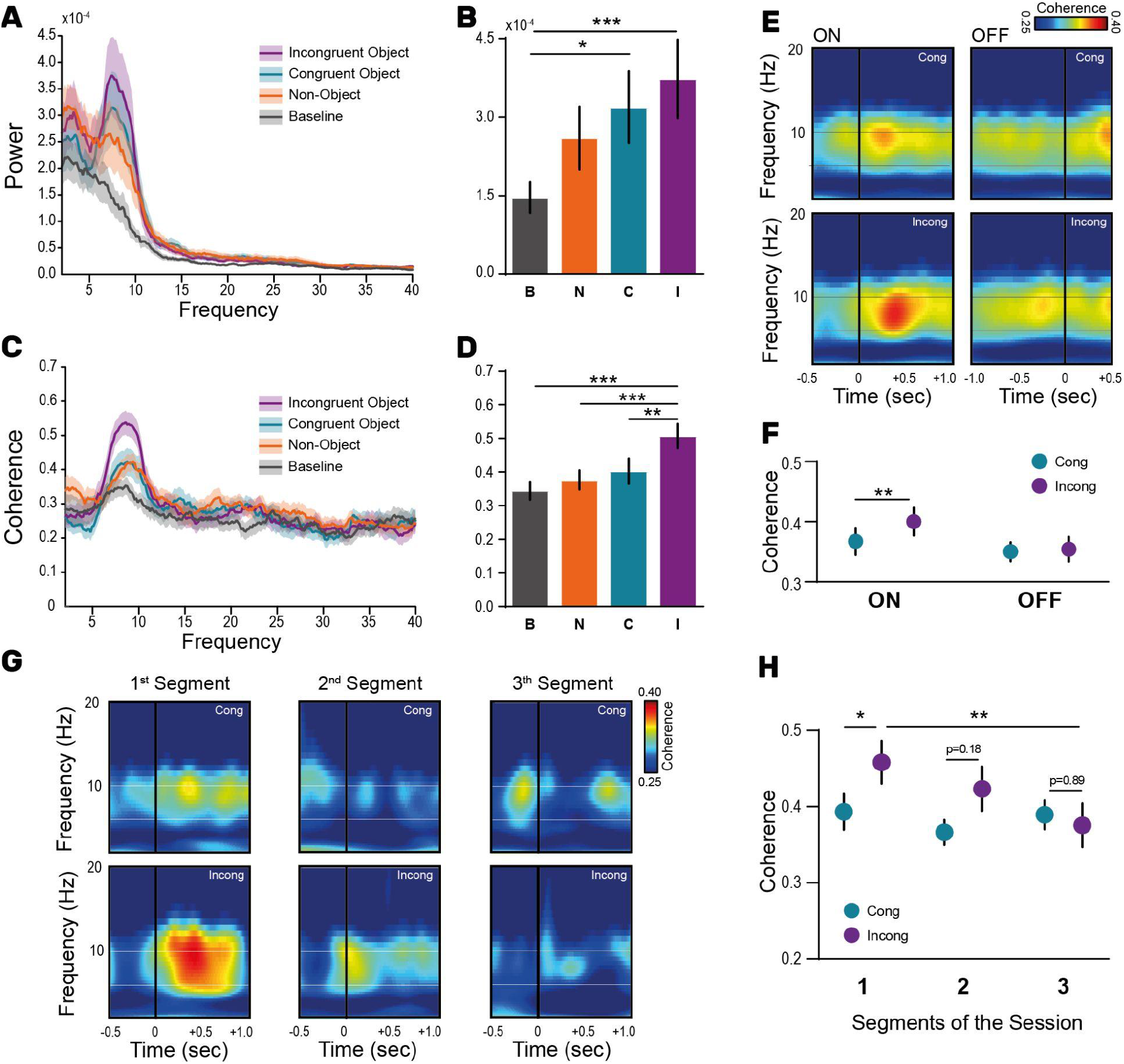
vHP-mPFC coherence is enhanced during incongruent object exploration. (A) vHPC-mPFC cross-spectrum analyzed during different behavioral events. Cross-power spectral density analysis was performed using LFP segments recorded during baseline condition (Grey) and during the test session separating Non-Object (Orange), Congruent Object (blue) and Incongruent Object (purple) events. (B) Quantification of theta power in the cross-spectrum for all conditions. One-way Repeated Measure ANOVA, N=21, ***p=0.0009, F=6.31. Bonferroni’s post-hoc test, ***p_I-B_=0.0005, *p_C-B_=0.035. (C) vHPC-mPFC coherence spectrum for all conditions. (D) Quantification of Coherence analysis. One-way Repeated Measure ANOVA, N=21, p<0.0001, F=11.46. Bonferroni’s post-hoc test, ***p_I-B_=0.0005, ***p_I-N_=0.0007, **p_I-C_=0.001. Bars represent mean ±SEM. (E) Peri-event coherence spectrograms centered at the beginning (ON, left) and at the end (OFF, right) of the congruent and incongruent object exploration events (upper and lower respectively). (F) Comparisons between coherence max value in the theta range during the beginning and the end of the congruent and incongruent events (blue and purple respectively). Two-way ANOVA, *p_Interaction_=0.03, F(1,20)=5.28. Bonferroni’s post-hoc test, **p=0.008, ***p=0.0003, ****p<0.0001. (G) Mean peri-event coherence spectrograms centered at the beginning of the congruent (upper) and incongruent (lower) exploration events across the test session of the OIC task. Object exploration events were split into three segments and the maximum coherence value was compared for both objects and across time (H). Two-way ANOVA, *p_interaction_=0.04, F(1.98,29,7)=3.53. Bonferroni’s post-hoc test, *p=0.023, **p=0.008.

**Figure 5:**
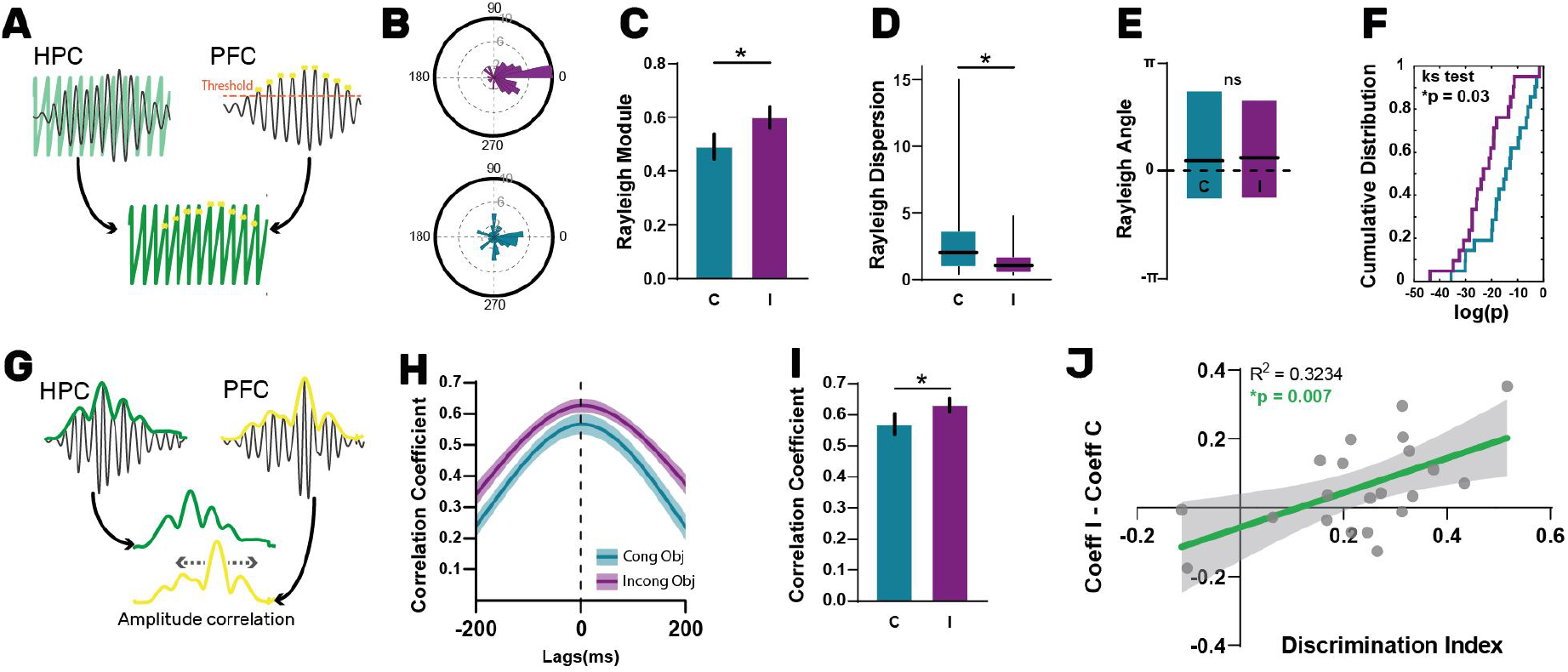
Synchronization of vHPC-mPFC theta oscillations is enhanced during exploration of the incongruent object and it correlates with the animals’ performance in the task. (A) Phase-locking analysis of vHPC-mPFC oscillations was performed by determining the vHPC phase at which mPFC theta peaks occur. (B) Example session showing the circular phase distributions during the exploration of the congruent (blue, module:0.1, dispersion: 14.77, p value:0.18) and incongruent objects (purple, module:0.56, dispersion: 1.06, p value:4.34×10^−9^). (C) Mean vector module obtained from the phase circular distribution of each test session during the exploration of the congruent (blue) and incongruent (purple) object. Paired t test, N = 21 per group, *p=0.017, t=2.59. Bars represent mean ± SEM. (D) Median Rayleigh’s vector dispersion for both conditions. Paired Wilcoxon test, N=21, *p=0.035. Graph represents median, 25-75 quartiles and range. (E) Angle of the mean vector obtained from circular phase distributions for congruent and incongruent conditions. Graph represents mean, 25-75 quartile. Wilcoxon paired test p=0.95. (F) Cumulative frequency function of Rayleigh’s test p values for the congruent (blue) and incongruent (purple) object. Two-sample Kolmogorov-Smirnov test, *p =0.03. (G) Scheme showing the method for calculating the cross-correlation of the instantaneous theta oscillation amplitude of the vHPC and the mPFC. (H) Mean correlation coefficient for congruent (blue line) and incongruent (purple line) conditions. Shaded areas represent ±SEM. (I) Correlation coefficient for both conditions at zero-lag. Paired t test, *p=0.04, t=2.09. Bars represent mean ± SEM. (J) Linear regression analysis between the behavioral discrimiation index and the difference of the correlation coefficient determined when animals explore the incongruent (Coeff I) and congruent object (Coeff C).

#### Phase Locking Analysis of LFP signals

mPFC and vHPC theta oscillations (6-10 Hz) during events of object exploration were isolated as described in the previous section. Here, the Hilbert transform was used to obtain the phase angle at every timestamp of the vHPC signal (“instantaneous phase”). mPFC theta oscillations peaks were detected and the instantaneous phase at which they occurred in the vHPC LFP recording was determined (**Figure 5A**) and plotted in as a circular histogram for each object condition (**Figure 5B**). Phase locking of mPFC theta peaks to hippocampal theta oscillation was determined by assessing deviation from uniformity in these circular plots using the Rayleigh test.

#### Tetrodes single-Units detection and clustering

Tetrode recordings from the mPFC were analyzed to extract single units as previously described in Alvarez et al., 2020. Spikes were detected, after high-pass filtering of the tetrodes signal (median filter, half window length: 2.4 ms) using NDManager (part of the published free-software package also including Klusters and NeuroScope, Hazan et al., 2006). Spikes were then assigned to single units (SU) by a semiautomatic spike sorting method based on spike’s relative amplitudes across four recording wires of a tetrode and PCA (Klusters software). Semiautomatic verification of clusters quality was performed using autocorrelograms, average spikes’ shape, and correlation matrices (Klusters) by a trained operator (**Figure 2C**, **Supp Table 1**). The stationarity of signals and the stability of unit isolation were visually verified for each recording session. Recording sessions in which cluster stability could not be confirmed were excluded from our dataset. Neurons were only included in our dataset if they met the criteria described elsewhere (Alvarez et al., 2020). All SU were divided into two subclasses (putative pyramidal cells and putative interneurons, **Figure 2D**) based on waveform shape as described previously (Barthó et al., 2004). In general, putative pyramidal cells showed relatively broad spike waveforms (with negative curvilinear shapes in the afterhyperpolarization phase), whereas putative interneurons had relatively narrow waveforms with positive curvilinear shapes (Alvarez et al 2020).

#### mPFC SU population analysis locked to object exploration

mPFC neurons firing rate was normalized (z-score) and the activity of all recorded cells was locked to the object exploration onset event. The z-score was calculated using +/- 20 seconds histograms centered on this event using a 0.2s bin. Neurons were then sorted according to their mean maximum activity in a +/- 0.2 seconds window around time zero at the perievent histogram (**Figure 6A**). mPFC Neurons were considered to respond to the congruent or incongruent object if they showed a change in activity in this time window greater than +/-1 SD.

**Figure 6:**
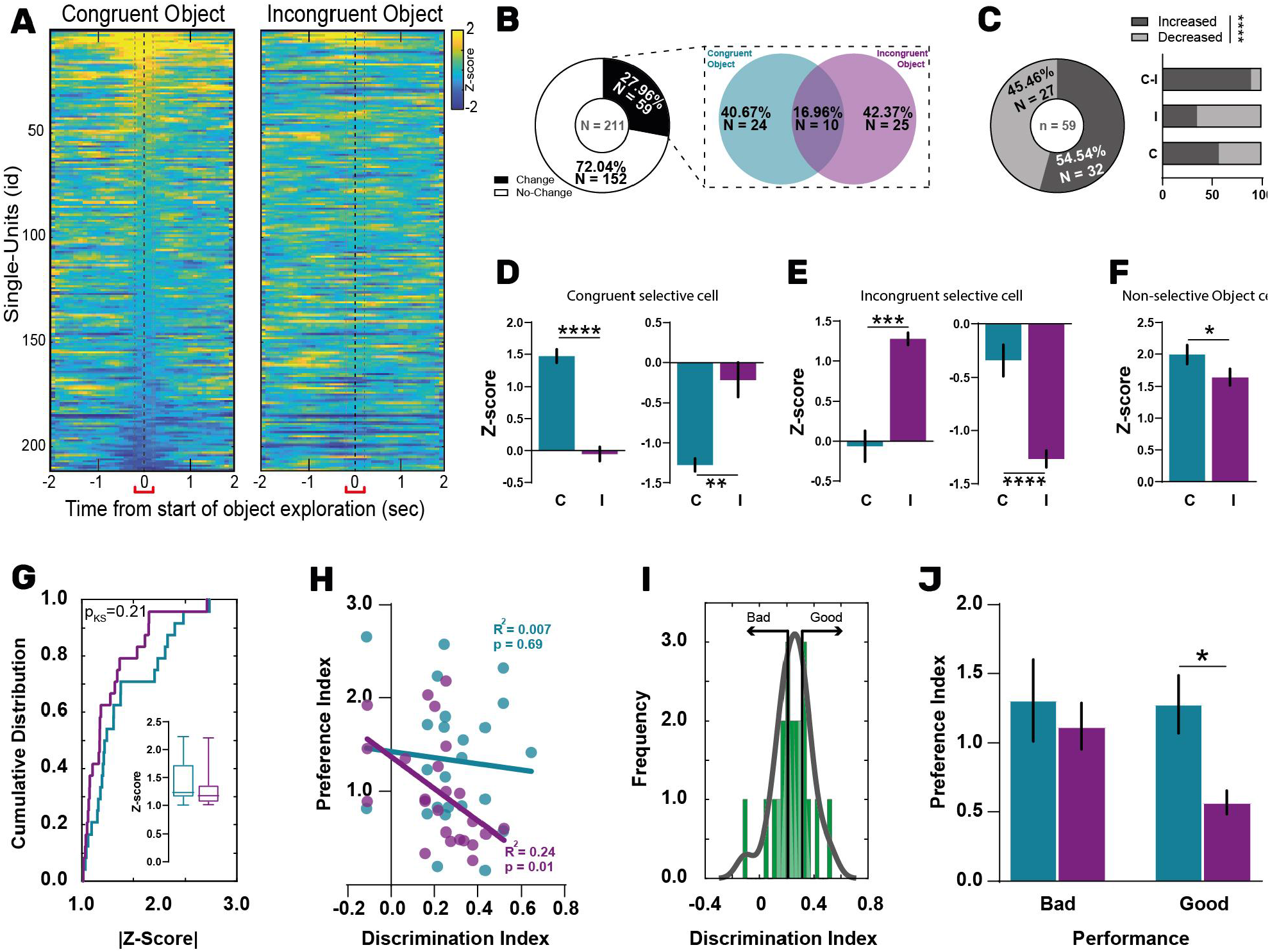
The incongruent object-context preference of mPFC neurons correlates with the animals’ performance during the resolution of the OIC task. (A) Peri-event time histograms locked to the beginning of the congruent (left) and incongruent (right) object exploration events for all single units recorded during the OIC recall. Single-units were sorted in descending order according to their mean z-score in a 400ms time window centered on the beginning of the congruent object exploration event. Then, the same sorting order was used to plot the activity of the neurons locked to the incongruent object event (right). (B) Percentage of mPFC neurons responding to the congruent (blue), incongruent (purple) and to both objects (intersection, non-selective). (C) Percentage of neuronal responses classified as excitations (dark grey) and inhibitions (light grey) according to their object preference (C:congruent; I: incongruent; C+I: non-selective; Chi-square=62.18, ****p<0.0001). (D) Average response of cells classified as congruent selective when animals explore the congruent (blue) and incongruent object (purple). Left: mean response of excitatory congruent cells responses (Paired t test, N=14, ****p<0.0001, t=9.36). Right: mean response of inhibitory responses (Paired t test, N=10, **p=0.0016, t=4.43). (E) Same as (D) for neurons classified as selective for the incongruent object(Left, Paired t test, N=9, ***p=0.0005, t=5.70; Right, Paired t test, N=16, ****p<0.0001, t=6.56). (F) Average Z-score for non-selective objects cells. Only cells responding with excitations were used (Paired t test, N=9, *p=0.039, t=2.46). (G) Cumulative frequency distribution of absolute z-score values for Congruent (blue line) and Incongruent (purple line) selective populations. Kolmogorov–Smirnov test, p=0.21. Inset: Median, 25-75 quartiles and range of the absolute z-score for both cell populations. Mann-Whitney test, *p=0.12, U=213. (H) Linear correlation between mPFC neuronal object preference index (|Z-score in Prefered condition - Z-score in Non-Prefered condition|) and the behavioral Discrimination Index. Two different linear regressions were calculated for the congruent (blue) and incongruent (purple) selective cells populations. (I) Frequency of behavioral discrimination indeces obtained for all test sessions. The distribution was splitted into two areas: i) left tail including 40% of sessions with smaller DI (Bad performance); ii) right tail including 40% of sessions with higher DI values (Good Performance). The envelope represents the probability density. (J) Neuronal preference index for congruent and incongruent selective cells in sessions where rats showed bad or good performance. Two-way ANOVA, *p_cell_=0.04, F(1,31)=4.61. Bonferroni’s post-hoc test, *p=0.04, t=2.36. Bars represent Mean ± SEM.

#### SU-LFP Phase Locking Analysis

The vHPC LFP signal acquired during object exploration events was filtered in the theta range as described in the sections above. A Hilbert transform was used to obtain the phase angle at every point of hippocampal theta oscillations. Next, the number of mPFC SU spikes occurring at different phase angles of the hippocampal theta rhythm was depicted in circular plots (**Supp Figure 5A**). Phase locking of mPFC SU discharges to hippocampal theta oscillations was determined by assessing deviation from uniformity in these circular plots with the Rayleigh test. To guarantee sufficient statistical power, only neurons that fired at least 25 spikes during the events of interest were considered for the phase-locking analysis.

### Postmortem histology

At the end of the experiments, rats were anesthetized and transcardially perfused with a cold saline solution followed by a buffered PFA solution (4% in PBS). Brains were removed, immersed overnight in the same fixative at 4°C. For cryopreservation, brains were immersed in different sucrose concentration solutions (10%, 20%, and 30%) for 24 hours each. Brains were maintained in sucrose 30% until further processed. Coronal brain sections were serially cut (40 µm) using a microtome with a freezing stage for histologic reconstructions. The location of the bipolar electrodes was assessed by visual examination of the mechanical tissue damage in the coronal sections stained with Safranin. Tetrodes localization was assessed by immersion of the electrodes in a red fluorescent dye (1,1′-dioctadecyl-3,3,3′,3′-tetramethylindocarbocyanine perchlorate, 100 mg/ml in acetone; DiI, Invitrogen) to detect the fluorescent material deposited in the tissue with an epifluorescence microscope (**Figure 2A**). Only rats whose electrodes were correctly placed were included in the analysis. One rat was excluded because of misplacement of the electrodes.

### Statistical Analysis

Statistical analyses were performed with GraphPad 8. Data were analyzed using two tailed unpaired Student’s t-tests when two groups were compared. For comparisons between two repeated-measured groups, a two-tail paired Student’s t-test was used. Paired One-way ANOVA followed by Bonferroni post-hoc test was performed for comparisons between more than two groups. If normality was not corroborated (using Shapiro-Wilk test), non-parametric test versions were performed (Man-Whitney for two groups or Kurskal-Wallis test for more than two groups). For correlations between continuous variables, we performed linear regressions. For distribution comparisons, a Two-sample Kolmogorov-Smirnov test was applied. In all cases, p values were considered to be statistically significant when p<0.05.

## Results

In rodents, episodic-like memories can be evaluated using object recognition paradigms based on the natural tendency of these animals to explore novelty. Particularly, the Object-in-Context (OIC) task is an object recognition task characterized by a strong contextual component (Morici et al., 2015) in which animals are trained to learn two different object-context associations. In this setting, both objects as well as the contexts are familiar to the animals. However, during the test phase, one of the objects is presented in a novel context generating a contextual incongruency (from now on Incongruent object, **Figure 1A**). Then, in order to solve the task, animals have to recognize a novel combination of previously known context and objects. Here, we used a slightly modified version of the OIC already published (see Methods section) to maximize the object exploration events for electrophysiological analysis of brain activity. Similar to what we have previously reported (Bekinschtein et al., 2013; Morici et al., 2018), animals showed increased exploratory time and a higher number of exploratory events for the incongruent object compared to the congruent one during the test phase of the task (**Figure 1 B-C**). Thus, the Discrimination Index (DI) was significantly different from zero (**Figure1D**). This preference for the incongruent object was observed and maintained across the entire test session (**Supp Figure 1B**). We also found a decrease in object exploration time across training sessions (**Supp Figure 1A**) suggesting equal habituation to the contexts and objects during the training phase. These results suggest that animals present a robust preference for the contextually mismatched object and that they can solve the task in this modified version (see M&M).

### vHPC-mPFC functional connectivity is required for the resolution of the OIC task

The vHPC and the mPFC appear as fundamental components of the circuit required to solve the OIC task (Morici et al., 2018). To establish if the interaction between vHPC-mPFC is required during the resolution of the novel OIC protocol we are using here, we performed a pharmacological disconnection experiment under these novel conditions. We trained cannulated rats in the OIC task (**Supp Figure 2**) and infused muscimol ipsi or contralaterally (IPSI or CONT, **Figure 1E** and **Supp Figure 2**) in the mPFC and vHPC before the test session. No differences in object exploration time for any of the conditions tested were observed (**Figure 1F** and **1G inset**) suggesting that muscimol infusions did not affect motor activity. However, CONT treatment significantly diminished the DI suggesting that bilateral interaction between vHPC and mPFC is necessary for the resolution of the task. To further analyze the type of connectivity between these structures we analyzed the ability to solve the OIC task in the IPSI treated group. Anatomical tracing studies suggest that ipsilateral is the most prominent interaction between these structures (Hoover and Vertes, 2007) suggesting that IPSI or CONT manipulation would have different effects on the OIC task. Supporting these results, we found that contrary to the CONT treated animals, the IPSI treatment did not affect the preference to explore the incongruent object (**Figure 1G**). This result can not be explained by differences in exploratory behavior during the training phase of the task since animales explored the objects similarly (**Supp Figure 2A**). Interestingly, we did not observe any differences in the DI when the same disconnection manipulation was performed during a novel-object recognition task (NOR), whose resolution does not depend on contextual information (**Supp Figure 3**). This suggests that this circuit is specifically recruited during contextually driven recognition memory recall. These results, together with what was described above, suggest that the protocol implemented here was effective to generate long lasting OIC memories that require a functional vHPC-mPFC communication during the retrieval phase.

### mPFC and vHPC theta oscillations increase during object exploration

Oscillations of neural circuits occur at a wide range of frequencies (Buzsáki and Draguhn, 2004) and help to filter, modulate and redirect information (Benchenane et al., 2010). LFP oscillatory signals participate in several processes going from cell ensembles formation to behavioral responses execution (Harris et al., 2003; Hasselmo, 2014; Hasselmo and Stern, 2014). Particularly, theta oscillations in the hippocampus as well as in the mPFC have been extensively associated with learning and memory. Thus, we decided to study the oscillatory profile of these structures and mPFC single unit activity during the recall of episodic-like-memories guided by contextual information. To do this, six male Wistar rats were implanted with four tetrodes in the mPFC and two stereodes in the vHPC (**Figure 2A**). The local field potential activity of these structures was recorded during the Test session of the Object-in-Context task (, **Figure 2B**). We also recorded a total of 211 single units from the mPFC (**Figure 2B** and **2C**). Isolated units were offline classified into two subclasses (putative pyramidal cells and interneurons) based on waveform and firing characteristics (**Figure 2D**). Putative pyramidal neurons were defined as cells with relatively broad spike waveforms and with negative curvilinear shapes in the afterhyperpolarization phase. Based on their relatively narrow waveform, 5% of all recorded cells were classified as putative interneurons, and most likely fast-spiking interneurons. Since putative interneurons were very few, all recorded mPFC cells were analyzed together.

To evaluate if theta oscillations were modulated during object exploration, a power-spectral density analysis of the LFP signal from the mPFC and the vHIP was performed on different behavioral periods. LFP activity was recorded prior to the beginning of the session, while the animals were still in their homecage, and it was used as a baseline to assess changes in LFP power during the task. Then, the test session was divided into periods of object exploration (Incongruent and Congruent) and time intervals where animals were not actively exploring the objects (non-object). We found that theta oscillations (6-10 Hz) were significantly enhanced in the mPFC and vHPC during object exploration periods (**Figure 3A** and **3C**). In the vHPC, we found that theta power was significantly increased during exploration of both objects compared to baseline and non-object periods (**Figure 3B**). In the mPFC, we observed an increase in theta oscillations during the exploration of the Incongruent object compared with the baseline and the non-object conditions (**Figure 3D**). In this structure, we did not observe statistical differences in theta power oscillation between the Incongruent and Congruent conditions (Bonferroni’s post-hoc test, p=0.415). These results indicate that theta oscillations are enhanced during active episodes of object exploration suggesting that they may be involved in the behaviors and cognitive processes associated with object recognition and retrieval.

### vHPC-mPFC LFP coordination shows high coherence in the theta range during the exploration of a contextually miss-matched object

It has been shown that the coordination between vHPC and mPFC in the theta band is enhanced in contextual-guided working memory (O’Neill et al., 2013; Liu et al., 2018) and rewarded episodic-like memory tasks (Place et al., 2016). Therefore, we addressed whether the functional connectivity between these structures changes dynamically during the test phase of the OIC task. We found a significant increase in the cross spectral power density in the theta range when animals were actively exploring either object compared to the baseline condition (**Figure 4A** and **4B**). Interestingly, vHPC-mPFC theta oscillations coherence was only increased when animals explore the contextually mismatched object, while the other conditions did not significantly differ from baseline (**Figure 4C** and **4D**). The same result was observed when we calculated the mean coherence values in the 6-10 Hz frequency range, and when we looked for the coherence value at the cross-spectrum peak (data not shown). Thus, while the mPFC and vHPC share an increase in theta activity irrespective of the explored object, they show a significant oscillatory synchronization only when the incongruent object is explored. To further address the dynamic modulation of this synchronization during object exploration, we built coherograms where coherence at different frequencies is plotted across time and centered at the beginning of the object exploration events (**Figure 4E** and **4G**). Interestingly, we found that the vHPC-mPFC coherence is higher predominantly at the beginning of the exploratory bout of the incongruent object (**Figure 4E** and **4F**). To further analyze the temporal dynamics of this process, we divided the test sessions in three segments and found that theta synchronization decreased throughout the session, specifically for the incongruent object (**Figure 4G** and **4H**). This suggests that an increase in vHPC-mPFC theta synchronization is required at the beginning of the session to recognize the object in context association as novel and to control the correct resolution of the task.

We performed two additional analyses assessing the phase synchronization between these structures. First, we performed a phase-locking analysis to determine how consistent the vHPC-mPFC theta oscillation phase was between structures. To do this, we looked at the vHPC phase at which mPFC theta oscillations peaks occurred (**Figure 5A**) and compared the phase circular distributions obtained when animals explored either object using the Rayleigh test (**Figure 5B**). We found that vHPC-mPFC theta oscillations show increased phase-locking in the incongruent condition compared with the congruent one. This was evidenced by a larger vector module (**Figure 5C**), and smaller dispersion (**Figure 5D**) and p-value (obtained when evaluating if the phases follow a uniform distribution, **Figure 5F**) of the circular phase distributions when animals explore the contextually mismatched object. We did not find significant differences between the angle of the phase distributions when comparing the congruent and incongruent condition (**Figure 5E**). In addition, we performed a cross correlation of instantaneous amplitudes of LFP theta oscillations (Adhikari et al., 2010). We found that the amplitude cross-correlation between vHPC-mPFC theta oscillations (**Figure 5G**) was significantly higher during the exploration of the incongruent object compared with the congruent one (**Figure 5H** and **5I**). This increase was not explained by an increase in theta amplitude per se, since we did not find a significant difference in theta amplitudes when animals explore the objects (**Supp Figure 4**). Interestingly, the difference between the cross-correlation value at zero-lag between the incongruent and congruent conditions positively correlated with the animals’ behavioral Discrimination Index (**Figure 5J**). This suggests that an increase in vHPC-mPFC synchronization during the exploration of the incongruent object is instrumental for the correct solving of the OIC task. Overall, the analysis of coherence, phase locking and instantaneous amplitude cross-correlation show a higher and more tight vHPC-mPFC synchronization during the exploration of the incongruent object than during the recognition of a known object-context match.

### mPFC single-units differentially encoding objects-in-context stimuli correlates with animals’ recall performance

Since vHPC-mPFC theta oscillations and functional connectivity are increased during object exploration, we addressed whether vHPC theta rhythms can modulate mPFC single-unit activity in the present task. To this aim, we recorded mPFC single-unit activity during the test phase of the OIC task. We found that 22,5% of mPFC recorded neurons (32/142) were significantly phase-locked to hippocampal theta rhythms (**Supp Figure 5A**). Among this neuron subpopulation, some units showed significant phase-locking when animals explore a particular object and some were non-selectively phased locked to object exploration (**Supp Figure 5B**). These results suggest that vHPC theta oscillations impact on the mPFC activity modulating the firing pattern.

Previous reports have shown that mPFC neuronal ensembles are modulated during working-memory (Spellman et al., 2015), and learning and memory tasks (Park et al., 2021). But it is still unclear if mPFC single-units can encode object-in-context information. To address this, we normalized the firing rate of individual mPFC neurons (z-score) and built peri-event time histograms (PETH) for every recorded cell centered on the beginning of each object exploration period (for Congruent and Incongruent objects). Neurons were then sorted according to their maximal response in a small time window around the moment animals begin exploring the objects (**Figure 6A**). Responses to object exploration were considered significant for cells showing changes in firing rate greater or smaller than mean ± 1SD (**Supp Figure 6A** and **B**). Overall, 59 of 211 (29%) neurons responded to object exploration (58 putative pyramidal cells and one putative fast spiking interneuron, **Figure 6B**). Once cells were sorted by their maximal response to both objects, two extra PETH were built to analyze how neurons showing significant responses to the one object responded to other one (e.g. **Figure 6A** shows the PETH of all neurons sorted by their maximal response to the congruent object and the same sorting order was used to evaluate how these cells respond to the incongruent object). We found that 24 and 25 out of 211 mPFC neurons (**Figure 6B**) showed a significant change in firing rate during the congruent and incongruent object exploration period respectively while 10 neurons responded to both of them (**Supp Figure 6, Figure 6C**). While the object selective neurons showed a very strong preference for either the congruent or incongruent object (**Figure 6D-E**), the non-selective object responding cells showed, on average, a slight preference for the congruent object (**Figure 6F**). Further analysis of object responses showed that most of the non-selective cells increased their activity during all object exploration events, while object selective neurons showed a more heterogeneous response profile of excitations and inhibitions (**Figure 6C**). To assess if the object selective responses were of similar amplitude irrespective of the congruent or incongruent nature of the context, we compared the cumulative frequency distributions of their response amplitudes and found no significant differences between them (**Figure 6G**). Altogether, these results suggest that distinct mPFC single unit subpopulations could differentially encode the representations of two familiar objects in context stimuli.

Finally, since we clustered distinct mPFC neuronal populations responding differently to the congruent and incongruent object, we asked if this neuronal object-in-context discrimination correlated with recall performance assessed with the DI. To do this, we evaluated how congruent and incongruent object selective cells responded to their preferred and non-preferred object by calculating a neuronal preference index. Interestingly, we found that the congruent-selective cells showed high object discrimination neural activity irrespective of the animals’ behavior, I. On the other hand, the neuronal object discrimination activity observed in the incongruent-selective cells significantly and negatively correlated to the behavioral discrimination index (**Figure 6H**). To further analyze this phenomenon, we compared the difference in the neuronal response to both objects for a subgroup of OIC tests where animals showed either good or bad performance in the task (**Figure 6I**). We found that when animals show a poor behavioral performance, both selective neuronal populations clearly discriminate between the contextually congruent and incongruent object with changes in firing rate (selectively increasing their firing for their prefered object). However, in those sessions where animals had a good behavioral performance and explored more the incongruent object, only the congruent cell subpopulation showed high neuronal preference index, while the incongruent cells fired similarly to both objects exploration (**Figure 6J**). Overall, strong congruent object selective responses do not inform about performance in the task. By contrast, a poor neuronal discrimination associated to a particular object correlates with a failure to recognize its association with the context. Together with the above findings these data suggest that lack of object discrimination at the neuronal level in the mPFC during enhanced theta synchronization with the vHPC might reflect the identification of a mismatch between the explored object/context pair and previously stored experiences.

## Discussion

This study examined the vHPC-mPFC coordination and mPFC activity during OIC retrieval. Our main findings are that: (1) vHPC-mPFC ipsilateral functional connectivity is required for the resolution of the OIC task; (2) theta oscillations power increases during the exploration of objects stimuli in the mPFC and vHPC; (3) the coherence, amplitude cross-correlation and phase-locking of theta oscillations between these structures increase significantly during the exploration of the contextually miss-matched object; (4) this increase in vHPC-mPFC synchronization is time restricted to the beginning of the test session and object exploratory bouts; (5) a greater vHPC-mPFC synchronization during the exploration of the contextually mismatched object correlates with the behavioral performance in the task; (6) there are subpopulations of putative pyramidal cells in the mPFC that selectively respond to object-context associations, and (7) their neuronal preference for one object correlates with the animals’ behavioral response during the resolution of the OIC task.

Episodic memory is the memory of events in their original spatio-temporal context: what happened, when and where it happened. Though the combination of these characteristics generates unique events, many times, specific features may be part of different memories. The mPFC is involved in memory processes mainly through strategic or cognitive control over retrieval processes. Eichenbaum and Preston proposed that the mPFC stores information during acquisition that is then used to flexibly select the memory trace to be retrieved, particularly when the animal is presented with conflictive information (Preston and Eichenbaum, 2013). The resolution of the OIC requires the recognition of the correct object context association when the animal is simultaneously presented with two object context pairs: one that has been experienced before (congruent object), while the other involves a new association between a familiar object with a context where it has not been presented before (incongruent object). In this scenario, two possible traces can be retrieved: the congruent memory appears as the most relevant to solve the task, while the incongruent object memory could interfere by activating a different episode (Eacott, 2004; Morici et al., 2015).

It has been proposed that in the OIC task, the mPFC uses contextual information to solve possible conflicting data arising from related memories. Behavioral studies suggest that this contextual information predominantly reaches the mPFC through ventral hippocampus monosynaptic ipsilateral projections (Preston & Eichenbaum, 2013, HOover & Vertes). Consistently with this theory, we found that vHPC-mPFC ipsilateral functional communication was not necessary to solve a novel object recognition task, but was instrumental for the resolution of the OIC.

Low frequency brain rhythms, like theta oscillations, are usually coherent over long distances and thought to link distributed cell assemblies. Thus, they have been proposed as attractors to control local and distant structures activity (Buzsáki, 2002; Stella and Treves, 2011). To evaluate the vHPC-mPFC functional connectivity in the OIC, we first studied the oscillatory activity present in both structures separately. Hippocampal theta oscillations are closely related to locomotion and animals’ speed, and they carry spatial information (O’Keefe & Recce, 1993; Buzsáki, 2002). Using an unrewarded task, like the OIC, we found that the power of ventral hippocampal theta oscillations increases during object exploration. Though an active behavior, object exploration does not require spatial shift, suggesting that hippocampal theta oscillations in this task, might be associated with the cognitive processes underlying the behavioral response. In the case of the PFC, studies performed in rodents, humans and non-human primates reported an increase of theta power during decision-making, attentional processes, and recently in a novel object recognition task (Liu et al., 2018, Tang et al., 2021; Wang et al., 2021). Here, we observed an increase in mPFC theta power during the exploration of the contextually incongruent object.

Since the vHPC and the mPFC showed increased theta power during object exploration, we analyzed the synchronization between both structures in the theta range during these events. Based on the anatomical connectivity and pharmacological experiments, we predicted that the resolution of the task would involve a tight communication between both structures. Increased theta vHPC-mPFC connectivity has been observed in spatial navigation, working memory (Sigurdsson et al., 2010; O’Neill et al., 2013), a novel object recognition task and rewarded object context associations (Alemany-González et al., 2020; Wang et al., 2021; Place et al., 2016). Consistently, we found that the vHPC and mPFC share an increase in theta activity irrespective of the explored object. Interestingly, further analysis demonstrated that the vHPC-mPFC theta synchronization increases significantly only when animals explore the incongruent object. This was evidenced by an increase in coherence, theta oscillation phase locking and amplitude correlation Interestingly, we also observed that vHPC-mPFC communication in the theta range impacts on the firing pattern of a proportion of mPFC neurons. This is evidenced by an increase in neuronal phase locking to this rhythm during object exploratory bouts.Altogether, this provides strong evidence for a differential requirement of vHPC-mPFC communication during the resolution of the task.

As we mentioned, the resolution of the OIC task requires the selection of the most relevant memory trace. This selection, as it happens in other tasks involving competing information, appears to involve the mPFC. Previous studies in humans (Kuhl et al., 2007, Wimber et al., 2015) and in rodents (Bekinschtein et al., 2018) have demonstrated that mPFC activity decreases as the cognitive control demands decline, suggesting that prefrontal interaction with downstreams structures is a dynamic and active process. Here, we observed that the vHPC-mPFC synchronization increases during the initial incongruent object exploration and then declines as object novelty decreases

Whatsmore, higher vHPC-mPFC synchronization during the exploration of the incongruent object, relative to the congruent one, correlates with the nature of the behavioral response (**Figure 5J**). The exploration of the incongruent object involves the recognition of the object itself (familiar to the animal) as well as the contextual mismatch. Therefore, the contextual information becomes key to solve the task. We hypothesize that the relative increase in vHPC-mPFC theta synchronization observed for the incongruent object, may help identify the object-context pair as novel and thus facilitate the correct selection of the relevant memory trace. This hypothesis is supported by the fact that the coherence between mPFC and vHPC is higher at the beginning of the test session, and declines as the object exploration progresses, and probably a new context-object linkage is created.

Several lesions and pharmacological experiments have shown that the mPFC is a key structure that participates in the retrieval of cue-guided object recognition paradigms (Morici et al 2015 for review). Due to the vast projections that arrive at the mPFC from associative regions (Klune et al., 2021), this region has been postulated as the orchestrator of memory reactivation in down-stream structures for different tasks (Liu et al., 2018, Tang et al., 2021; Wang et al., 2021, eichenbaum 2017). We have previously shown that activation of the vHPC as well as its interaction with the mPFC during the resolution of the OIC affects the labilization of OIC memory traces in the Perirhinal Cortex (Morici et al., 2018). Interestingly, we found that it is the memory trace of the congruent object the one that normally was reactivated, supporting that it is the most relevant trace to solve the task. Also, when both memory traces were reactivated, animals showed a bad behavioral performance in the test. Then, it is relevant to analyze the cellular signature within the mPFC during the resolution of the task. We identified two neuronal subpopulations that responded selectively to the congruent and incongruent objects. As in many other behavioral experiments, we found some variability between trials in the animals’ performance in the task. This normal variability gave us the opportunity to analyze if the behavioral performance correlated with the mPFC neuronal activity. We found that the selective congruent object neurons always had greater responses when exploring their preferred object (i.e. high preference index) regardless of the animals’ behavioral performance. In contrast, the selective incongruent object subpopulation showed neuronal preference indexes that correlated with the behavioral performance in the task. When animals had a poor behavioral DI, the incongruent object subpopulation showed high neuronal discrimination activity, similarly to the congruent object subpopulation. However, when the behavioral DI was high, the incongruent neuronal subpopulation responded similarly to both objects. (i.e low preference index).

As we mentioned, the mPFC is thought to control the retrieval of the most relevant memory trace in downstream structures. But, how may the mPFC direct the reactivation of different memory traces? We hypothesize that the activation of specific neuronal ensembles with high preference for a particular object-context pair could drive the retrieval of the original memory trace for that match. In the context of the present task: i) when both neuronal subpopulations show high preference, the incongruent object would be retrieving an incorrect trace, related to a past experience of this object in another context and leading to a poor object-in-context recognition; ii) when the incongruent object subpopulation shows low preference index, only the most relevant memory trace is retrieved. We interpret that this low preference index of the incronguent object ensemble generates a noisy signaling within the mPFC that helps inhibit the retrieval of the less relevant memory trace. Human studies have shown that prefrontal regions are engaged during retrieval inhibition and that they modulate the activity of downstream areas (Anderson and Floresco, 2021). Interestingly, functional homologous regions in rodents, like the mPFC, have also been associated with retrieval inhibition (Bekinschtein et al., 2018). What are the local processes that enable the cortex to control retrieval interferenceThough our work does not answer this question, it does provide some interesting observations. We identified two distinct and object-context selective neuronal ensembles whose activity correlated with the behavioral response. It has been suggested that mPFC ensembles select the most appropriate behavioral response by acting as comparators (McKenzie et al., 2016) between the present information and that acquired during training. When animals show good performance, the lack of contextual coincidence is reflected in the incongruent neuronal subpopulation as a decrease in the signal to noise ratio (i.e., a small difference between the signals emitted during exploration of the object-context pairs) that drives the inhibition of the incongruent memory trace. What type of information drives the change of ratio observed? Our results do not allow us to answer this question. It could come from a local circuit controlling the firing pattern of mPFC cells or, as mentioned above, it could be driven by other structures like the vHPC that send contextual information to the mPFC relevant to solve the task. For instance, the contextual inconsistency might be coded as an increase in vHP-mPFC synchronization during the incongruent object exploration and might be read by mPFC neurons to compare recent and old information.

In summary, we found that the resolution of the OIC task requires the functional connectivity between the VHPC and mPFC. This connectivity is reflected as an increased synchronization between both structures, especially when the animals explore the incongruent object, suggesting that the contextual information is required by the mPFC to select the correct memory trace. We found two mPFC neuronal ensembles that might drive the retrieval during the resolution of the OIC task. The inhibition of the less relevant trace appears to be achieved by a decrease in the signal to noise ratio in the ensemble responding to the incongruent object.

## Acknowledgments

We would like to thank Dr Juan Belforte, Dr. Pedro Bekinschtein and Dr Gustavo Murer for their critical reading and comments on the manuscript. We would also like to thank Graciela Ortega, Veronica Risso and Jessica Unger for technical support.

