## Supplementary Figures for "Hippocampal-Prefrontal cortex network dynamics predict performance during retrieval in a context-guided object memory task"

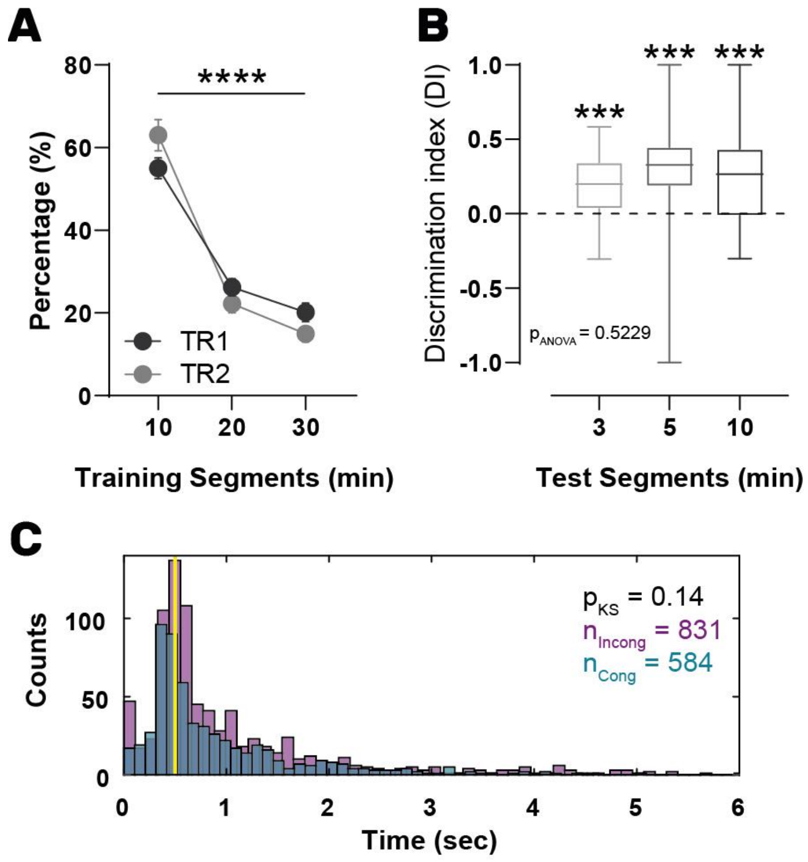


**Supplementary Figure 1: Behavioral data of the OIC task in implanted rats**. (A) Percentage of object exploration time across training sessions. Two-way ANOVA, N=21 per group, $p_{Interaction}$=0.13, $F_{Interaction}$=2.08,****$p_{Time}$<0.0001,$F_{Time}$=89.58. Each data point represents mean ±SEM, (B) Discrimination Index acoss time during test sessions. Friedman test, N=21 per bar, p=0.23, F=2.95. Graphs represent median, 25-75 quartiles and range. (C) Histograms of object exploration events durantios (blue bars: congruent; purple bars: incongruent). Two-sample Kolmogorov-Smirnov test, p=0.14. The yellow line is placed on 500ms, -the minimum event duration used in the LFP analysis-.

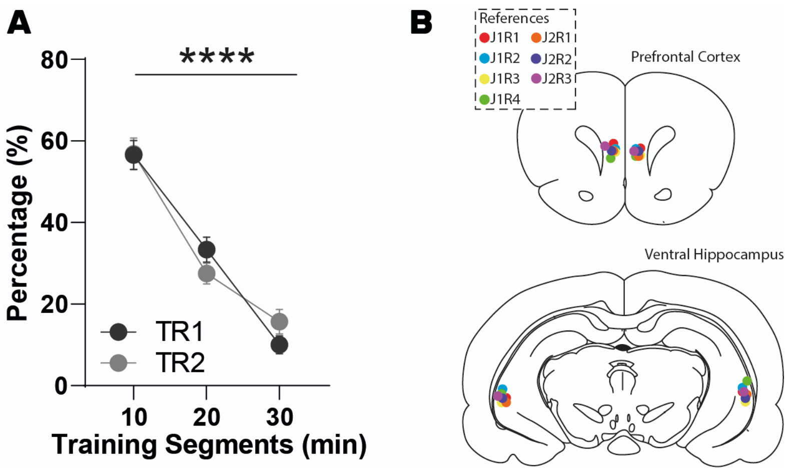


**Supplementary Figure 2: Behavioral data from the pharmacological disconnection experiment during the recall of the OIC task**. (A) Percentage of object exploration time across the training sessions. Two-way ANOVA, N=14 sessions per condition, $p_{Interaction}$=0.32, F=1.17, ****$p_{Time}$<0.0001, $F_{Time}$=68.16. Plotted Mean ±SEM. (B) Schematic representation of cannula placement in each rat. Upper: Prefrontal Cortex, Lower: Ventral Hippocampus.

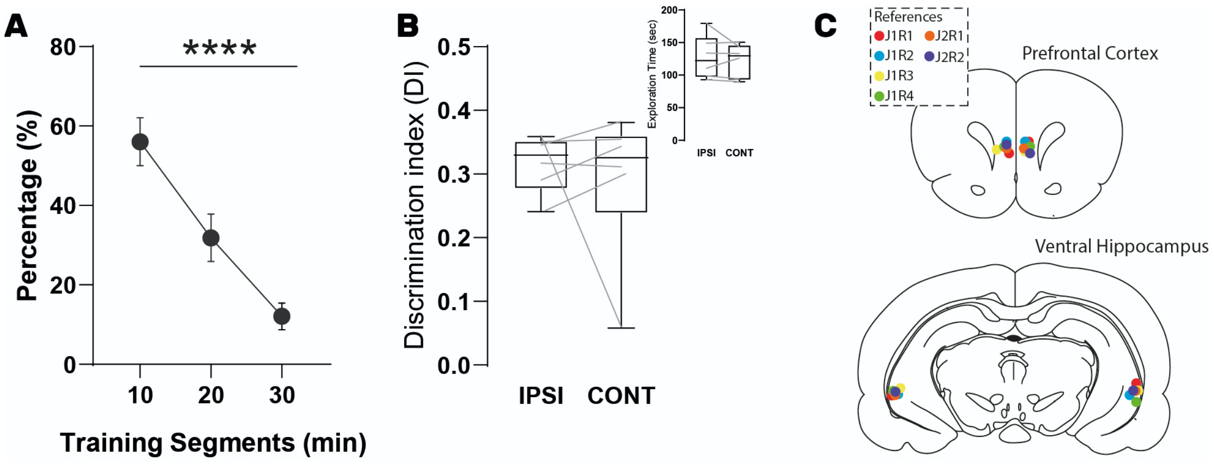


**Supplementary Figure 3: Behavioral data from the pharmacological disconnection experiment performed during the recall of the NOR task**. (A) Percentage of object exploration time across the training sessions. One-way ANOVA, N=12 sessions per condition, ****p<0.0001, F=140.4. Graphs represent mean ±SEM. (B) Discrimination Index for ipsilateral (IPSI) and contralateral (CONT) conditions. Each rat received both treatments in a pseudorandomly manner. $DI=(T_{Novel}-T_{Known})/(T_{Novel}+T_{Known})$. Wilcoxon test, p=0.69. inset: total object exploration time observed during the test session. Paired t test, p=0.54, t=0.66. Graphs represent median, 25-75 quartiles and range. (C) Schematic representation of cannula placement in each rat from this experiment. Upper: Prefrontal Cortex, Lower: Ventral Hippocampus.

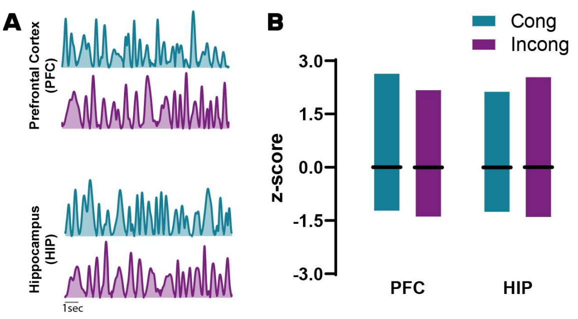


**Supplementary Figure 4: mPFC and vHP theta oscillation instantaneous amplitude**. (A) Example traces of mPFC (upper) and vHPC (lower) theta oscillation amplitudes for the congruent (blue) and incongruent (purple) conditions . B) Quantification of the area under the curve for each structure for both conditions and structures. Graphs represent mean, and range. Two-way ANOVA, $p_{interaction}>0.99$, $F(1,40)=1.2e^{-029}$.

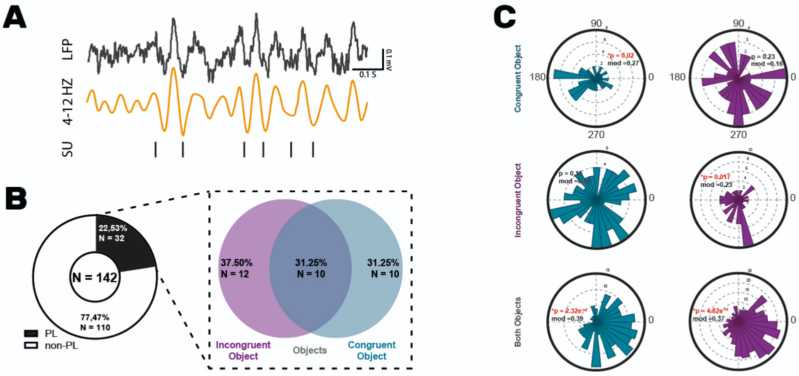


**Supplementary Figure 5: Prefrontal cortex single-units phase locking analysis to hippocampal theta rhythm**. (A) Example of the general procedure for this analysis. Hippocampal LFP signal was filtered (4-12 Hz, orange trace) and a Hilbert transform was used to obtain the instantaneous phase of vHPC theta oscillations at which mPFC action potentials (vertical bars) occurred. . (B) Percentage of mPFC single-units significantly phased locked to the vHPC theta rhythm. Inset, proportions of cells locked during exploration of one or both objects. (C) Examples of circular distributions of phases for cells locked during exploration of the congruent (upper), incongruent (middel) or both (lower) objects.

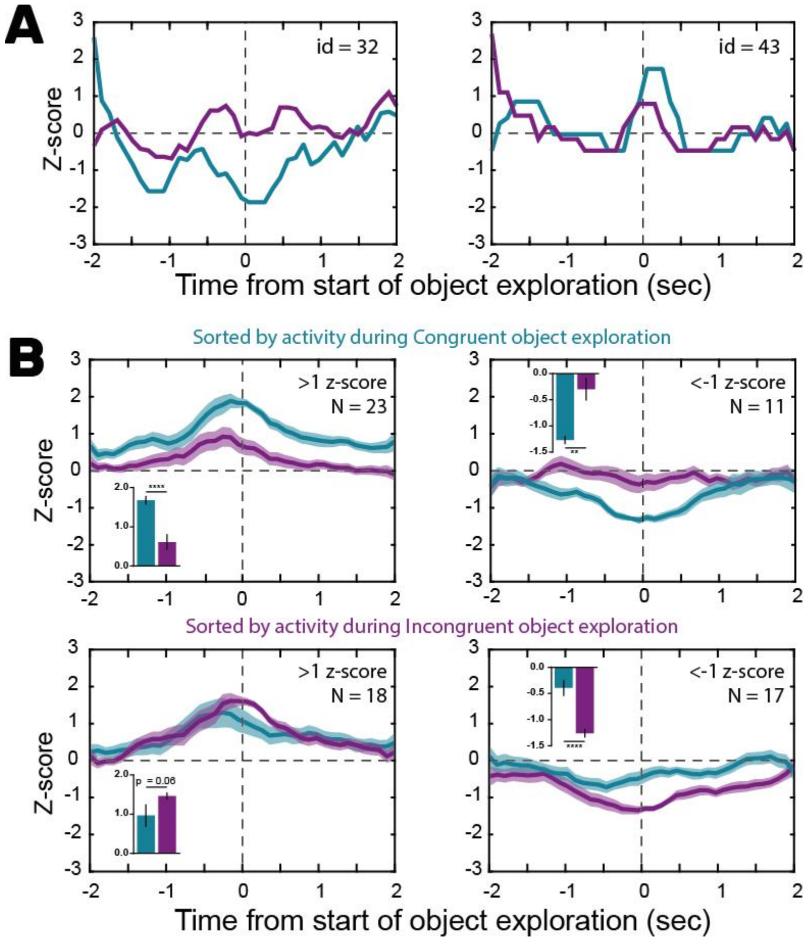


**Supplementary Figure 6: Single-Units classification according to their response during objects exploration events.** (A) Examples of two cells responding with an inhibition (left) or excitation (right) during the exploration of the congruent object. (B) Mean z-score for neurons showing excitations (left) and inhibitions (right) during the exploration of the congruent (upper) or incongruent (lower) object. Insets represent the mean z-score value in a time window of 400ms centered at time 0 for all conditions. Bars represent mean ± SEM. Paired t test, >1 in Congruent, ***p<0.0001, t=6.43, <-1 in Congruent, **p=0.002, t=4.11, >1 in Incongruent, p=0.06, t=2.02, <-1 in Incongruent, ****p<0.0001, t=6.02.

| Rat id | # recorded sessions | # Object Exploration Events | | # Single-Units | |
|---|---|---|---|---|---|
| | | Congruent | Incongruent | Pyr | FS |
| 3 | 1 | 39 | 49 | 12 | 1 |
| 4 | 4 | 168 | 233 | 16 | 1 |
| 5 | 4 | 116 | 141 | 1 | |
| 6 | 6 | 126 | 228 | 53 | 7 |
| 7 | 4 | 46 | 85 | 21 | 2 |
| 9 | 7 | 226 | 268 | 97 | 0 |

**Supplementary Table 1: Summary of the data obtained for each animal during the test session of the electrophysiological experiments.** . We report the number of sessions recorded per rat, the number of object exploration events registered per animal, and the amount of putative pyramidal (Pyr) and fast spiking interneurons (FS) recorded in each rat.
